# Normalization of MRI T1w between-scan effects for improved longitudinal volumetric estimates

**DOI:** 10.1101/2022.06.19.496756

**Authors:** Donatas Sederevičius, Atle Bjørnerud, Kristine B. Walhovd, Anders M. Fjell

## Abstract

Variations in image intensities and contrasts between magnetic resonance imaging (MRI) acquisitions affect the subsequent image processing and its derived outcomes. Therefore, comparability between acquisitions is improved if we reduce these variations. This is especially relevant for longitudinal studies where a change of scanner or acquisition protocol often happens between subsequent examinations. In this study, we use a robust intensity distribution alignment (RIDA) method to reduce between-scan effects and improve longitudinal volume change estimates between two MRI scanners – Siemens 1.5T Avanto and 3T Skyra. The method is based on MRI T1w images acquired in close succession and robustly aligns two cumulative distribution functions of voxel intensities to harmonize image intensities and improve image-derived outcomes of a range of subcortical brain. We compare RIDA with volume-based correction - a simple linear regression model. In both cases, we derive intensity and volume transformations from a training dataset of 20 participants scanned on both scanners on the same day and apply to an independent longitudinal test dataset of 243 participants. All participants in the test set were scanned at the Avanto scanner at the baseline and then at the Avanto and Skyra scanners on the same day at the follow-up, on average 4.4 years (sd = 0.5 years) later. This allowed us to directly assess the effect of scanner and protocol change on the longitudinal change estimates. Eight subcortical brain regions were segmented using SAMSEG, and annualized symmetrized percent change in volume between time points was calculated. We find that RIDA significantly reduces between-scan effects and improves longitudinal volume estimates for the amygdala and lateral ventricles. It also reduces between-scan effects for caudate, putamen, and thalamus, but not as much as linear regression models. Whether the method will be useful for a particular study will depend on the image intensity profiles of the scans. Therefore, a pilot study of double-scanned participants is recommended to assess the advantages of the method for the analysis in question.

## 1. Introduction

The development of techniques for quantifying changes in brain structures by using magnetic resonance imaging (MRI) has accelerated over the past decade. We can now measure subtle changes in, for instance, the thickness of the cerebral cortex or volume of subcortical structures (Holland et al., 2009). This development has enormous clinical potential for several patient groups, especially slow-progressing age-related disorders such as Alzheimer’s disease (AD) (Jack et al., 2015). It has been shown that quantitative MRI can be used to classify AD patients and age-matched controls with high accuracy (Chepkoech et al., 2016; Fjell et al., 2010). Of most interest, disease progression can be monitored prior to the stage of clinical diagnosis (Jack et al., 2013, 2010) and before the disease has progressed so far that any attempt at treatment will be futile due to the extensive brain damage. Thus, quantitative MRI can detect and monitor brain changes at a stage when only subtle clinical symptoms are seen, which is also critical for differential diagnosis.

Despite these advantages, quantitative brain MRI is not commonly used in clinical settings to monitor the rate of atrophy over time. A significant obstacle is that the results are distorted by subtle differences between scanners, protocols, and head coils, affecting image contrast and quality. This makes quantitative values challenging to compare to norm materials and previous examinations of the same patient.

This work is a continuation of our previous work (Sederevicius et al., 2022), in which a robust intensity distribution alignment (RIDA) method is presented. There we have developed an image harmonization by the RIDA, which we contrasted with the other two methods: multisite image harmonization by cumulative distribution function alignment (*mica*) (Wrobel et al., 2020) and Removal of Artificial Voxel Effect by Linear regression (RAVEL) (Fortin et al., 2016). The RIDA significantly improved consistency in volumetric measurements between two different MRI T1w images acquired on the same day for six studies. In this work, we apply RIDA to a longitudinal dataset to assess whether the improved between-image correspondence could also improve the longitudinal change estimations in studies where the scanner and protocol are replaced between the baseline and follow-up.

We use two distinct between-scan effects normalization approaches: volume versus intensity corrections. For the volume correction, we use linear regression models on the image-derived volumetric estimates of the brain structures and estimate volume transformations between Siemens 1.5T Avanto and 3T Skyra images. For the intensity correction, we deal with the raw image intensities mapping intensity profile from one scanner to another using RIDA. We use a fully automated longitudinal whole-brain segmentation method called SAMSEG (Puonti et al., 2016) (part of FreeSurfer v7.1) to label eight brain structures of interest: amygdala, caudate, hippocampus, lateral ventricle, inferior lateral ventricle, pallidum, putamen, and thalamus.

This work aims to address the issue of between-scan effects utilizing the two normalization approaches in the context of a longitudinal study. Here we assumed that between-scan effects were due to the change of scanner and T1w acquisition protocol parameters. We used the previously developed RIDA method to assess whether longitudinal change estimates could be improved. We contrasted these results to the effect of using a simple linear regression model. In both cases, we derived intensity and volume transformations from a training dataset of 20 participants scanned on an Avanto 1.5T and a Skyra 3T scanner on the same day and applied this to a longitudinal test dataset of two visits for 243 subjects. In the test dataset, all participants were scanned at the Avanto scanner at baseline and both the Avanto and the Skyra scanner at the follow-up. This allowed us to compare the methods to correct the scanner and protocol change to be an ideal situation of the same scanner being used at both visits.

## 2. Material and methods

### 2.1. Datasets

In this work, we used two datasets: training and test. The training set consisted of 20 cognitively healthy participants (18 females, age range from 20 to 36 years, mean age = 27.3 years, sd = 4.5 years), each scanned on the same day using two models of Siemens MRI scanners (Siemens Medical Solutions, Erlangen, Germany): 1.5T Avanto and 3T Skyra, at Rikshospitalet, Oslo University Hospital. We used the following T1w acquisition parameters: TR = 2400 ms, TE = 3.61 ms, TI = 1000ms, FA = 8°, matrix = 240×240×160, voxel size = 1.25×1.25×1.2 mm^3^ for the Avanto, and TR = 2300 ms, TE = 2.98 ms, TI = 850 ms, FA = 8°, matrix = 256×256×176, voxel size = 1×1×1 mm^3^ for the Skyra. The purpose of the training dataset was to derive image harmonization transformations which could be applied to the test dataset.

The test set was independent of the training set and consisted of 243 cognitively healthy participants (136 females, age range from 10 to 81 years) scanned longitudinally twice (mean years between two scans = 4.4 years, sd = 0.5. years). At visit 1, every participant was scanned with the Avanto scanner; at visit 2, every participant was scanned with both the Avanto and Skyra on the same day. The same MRI T1w acquisition parameters were used for both scanners as in the training dataset.

### 2.2. MRI pre-processing

We pre-processed both datasets in the same way as described in (Sederevicius et al., 2022). Briefly, the images of each participant were corrected for geometrical variations due to the scanner’s gradient non-linearities (Jovicich et al., 2006), resampled to 256×256×256 dimensions and 1×1×1 mm^3^ voxel size, corrected for intensity inhomogeneities (Puonti et al., 2016) and White Stripe normalized (Shinohara et al., 2014).

### 2.3. Methods

We used two fundamentally distinct approaches to address the issue of between-scan effects. For the first approach, an intensity correction, we used a robust intensity distribution alignment (RIDA) method to map an intensity profile of an image resulting from the Skyra scanner to an intensity profile of an image from the Avanto scanner (Sederevicius et al., 2022). Briefly, we calculated the cumulative distribution functions (CDFs) of the two images’ subcortical intensity values (as described in the original paper). We mapped the Skyra intensity distribution to the Avanto intensity distribution using a simple linear interpolation between two intensity profiles (CDFs). We derived an intensity mapping for each participant in the training data set and averaged it to come up with the final intensity transformation used to transform Skyra-like to Avanto-like image intensities in the test dataset. Therefore, each subject had an additional image at visit 2, Avanto-predicted. Finally, we processed all images of both visits with longitudinal SAMSEG (Cerri et al., 2020; Iglesias et al., 2016) to obtain volumetric measurements of eight subcortical brain structures of interest: amygdala, caudate, hippocampus, lateral ventricle, inferior lateral ventricle, pallidum, putamen, and thalamus.

For the second approach, a volume correction, we used a simple linear regression (LR) model to estimate a linear relationship between volumetric measurements resulting from Skyra (*V*_*s*_) and Avanto (*V*_*A*_) images:

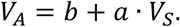

The coefficients *a* and *b* were estimated from the training data set. We applied the regression model to each brain structure of interest, resulting in eight different models. The inherent difference between the two approaches is that an intensity transformation changes raw image intensities, then image-derived measures are obtained, whereas volume correction operates on image-derived measures. In both cases, however, we use the training dataset to derive both transformations – the intensity and volume.

We analyzed within-visit volume differences (at timepoint 2) and between-visit longitudinal volume changes. For the within-visit, we looked at the following volume differences of visit 2: 1) Avanto vs. Skyra, 2) Avanto vs. Skyra volume-corrected (VC), and 3) Avanto vs. Skyra intensity-corrected (IC). The Avanto vs. Skyra represented volume differences influenced by the between-scan effects, whereas 2) and 3) represented volume differences after the scan effects were reduced by the volume and intensity corrections, respectively.

For the between-visit, we quantified the following longitudinal volume changes: 1) visit 1 Avanto and visit 2 Avanto (Avanto-Avanto), 2) visit 1 Avanto and visit 2 Skyra (Avanto-Skyra), 3) visit 1 Avanto and visit 2 Skyra volume-corrected (Avanto-Skyra VC) and 4) visit 1 Avanto and visit 2 Skyra intensity-corrected (Avanto-Skyra IC). The Avanto-Avanto reflects a scenario as if we had longitudinal data of the same scanner and MRI T1w protocol and stands as reference measurements. At the same time, the Avanto-Skyra represents a typical situation where we have scan effects due to the change of a scanner/protocol that often happens in longitudinal studies. The latter two, Avanto-Skyra VC and Avanto-Skyra IC, are longitudinal estimates between two visits after between-scan effects normalization, namely volume and intensity corrections.

### 2.4. Statistics

We split the longitudinal test dataset into three age groups: 1) less than 20 years (91 participants, 44 females, mean years between visits = 4.4 years, sd = 0.4 years), 2) between 20 and 40 years (43 participants, 25 females, mean years between visits = 4.6 years, sd = 0.5 years), and 3) greater than 40 years (109 participants, 67 females, mean years between visits = 4.3 years, sd = 0.5 years). The second group had a similar age range to the training dataset, so we called it a *representative* group. We referred to the first and the last group as *development* and aging, respectively. We used all three groups to report longitudinal volume change between two visits before and after the scan-effect normalization.

To calculate within-visit volume differences before and after between-scan effects normalization, we used absolute symmetrized percentage difference (ASPD):

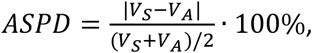

where *V*_*A*_ and *V*_*S*_ are estimated brain volumes at visit 2 for the Avanto and Skyra images, respectively.

To calculate the volume change from visit 1 to 2 and to evaluate the performance of both scan-effect normalization methods, we used annualized symmetrized percentage change (ASPC):

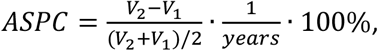

where *V*_1_ and *V*_2_ are estimated brain structure volumes at visits 1 and 2, respectively, and *years* is the time between MRI visits expressed in years. We calculated the mean ASPC for each structure (averaged between left and right hemispheres) among participants and tested the statistical significance of between-scan effects normalization using paired samples t-test.

All statistical analyses described above were done using R statistical software package v3.6.3 (R Core Team, 2020) and its packages: *ggplot2* (Wickham, 2016), *ggpubr* (Kassambara, 2020), and *dplyr* (Wickham et al., 2020).

## 3. Results and Discussion

### 3.1. Volume and intensity corrections

Fig. 1 and Fig. 2 present volume and intensity correction transformations derived from the training dataset of 20 participants. The linear models were used to convert longitudinal SAMSEG volumetric estimates of the Skyra images to the Avanto-based estimates. All slope coefficients were close to 1, meaning that the differences between the two types of images were mainly explained by the offset - the difference between mean volumes of the Avanto and Skyra. For the intensity-based correction, the individual intensity profiles were close to the mean intensity profile, meaning there was little variance between the participants.

**Fig. 1.**
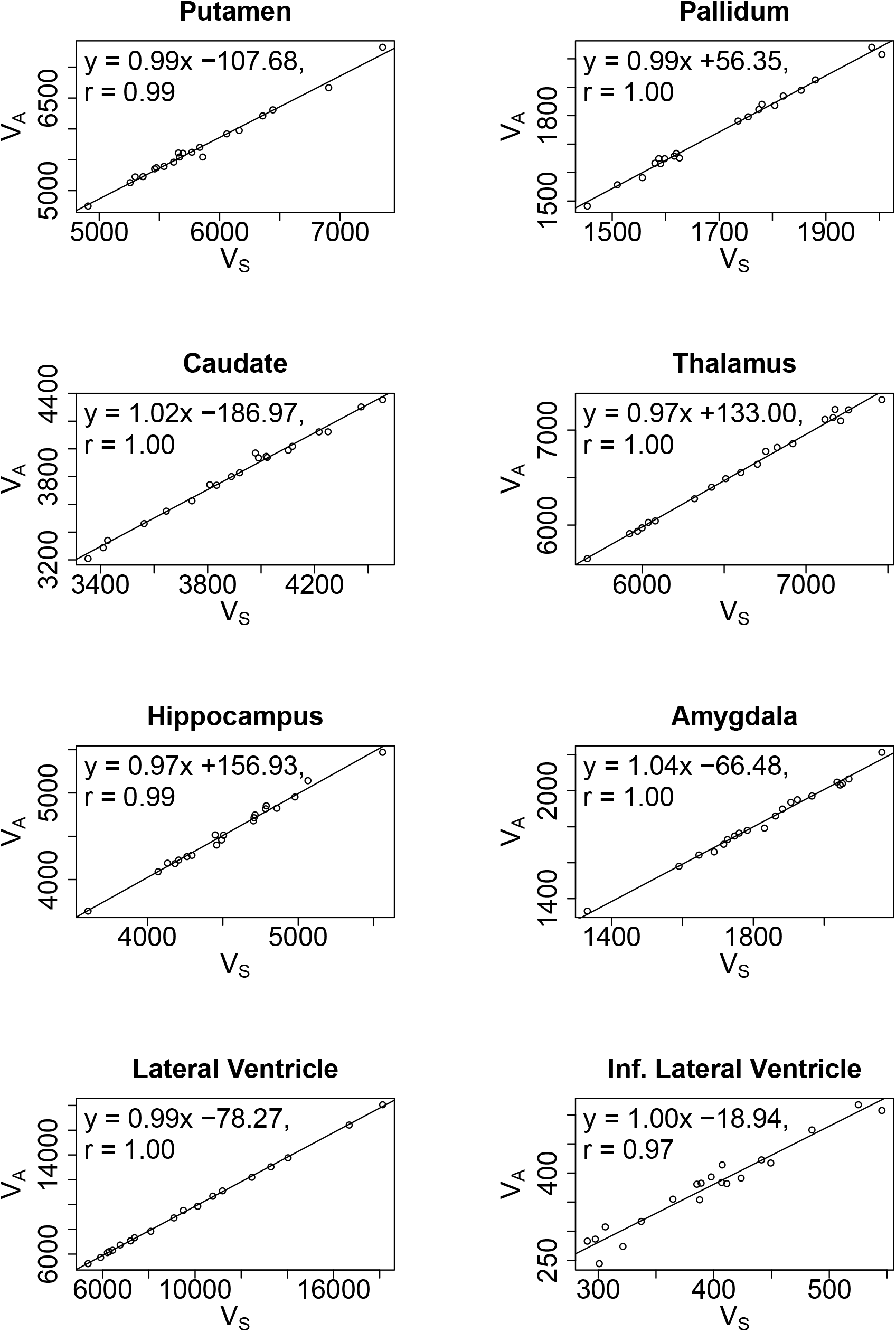
Linear models derived from the training dataset of 20 participants scanned on the same day with the Avanto and |Skyra scanners. SAMSEG estimated Skyra-like volumes (V_S_) are converted to Avanto-like (V_A_) volumes. The letter ‘r’ indicates the Pearson correlation coefficient below the linear model.

**Fig. 2.**
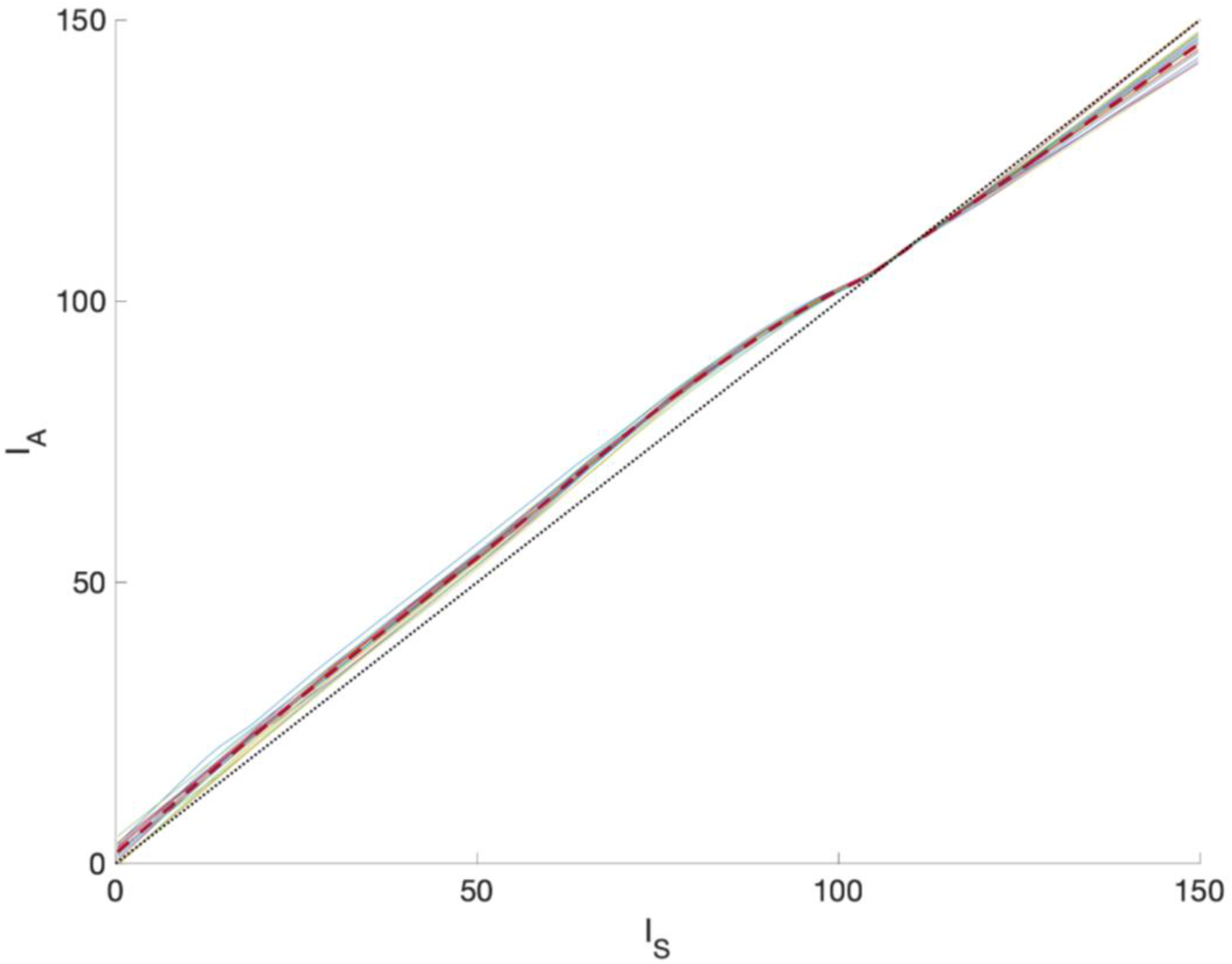
Individual and mean intensity mappings of the training dataset. Intensity values of the Skyra image (I_S_) are converted to the Avanto-like intensities (I_A_). The solid lines show the individual intensity profiles of 20 subjects, and the red dashed line shows the mean of all. The identity transformation is shown by the black dashed straight line.

### 3.2. Within-visit volume differences

Fig. 3 presents within-visit volume differences before (RAW) and after between-scan effects normalization (RIDA and LR) for the selected subcortical structures of the 243 test participants. RIDA yielded lower volume ASPD values for the amygdala, caudate, lateral ventricles, putamen, and thalamus. However, caudate, putamen, and thalamus volume corrections by the linear regression resulted in significantly lower ASPD values than RIDA (p < 0.05). Neither of the correction methods managed to improve the volumetric differences for the hippocampus and pallidum. Paired samples t-test with Bonferroni multiple comparisons correction indicated statistically significant (p < 0.05) differences between correction methods except for hippocampus (RIDA vs. LR) and pallidum (RAW vs. RIDA). The numerical results of Fig. 3 are presented in the appendix, Table A.1.

**Fig. 3.**
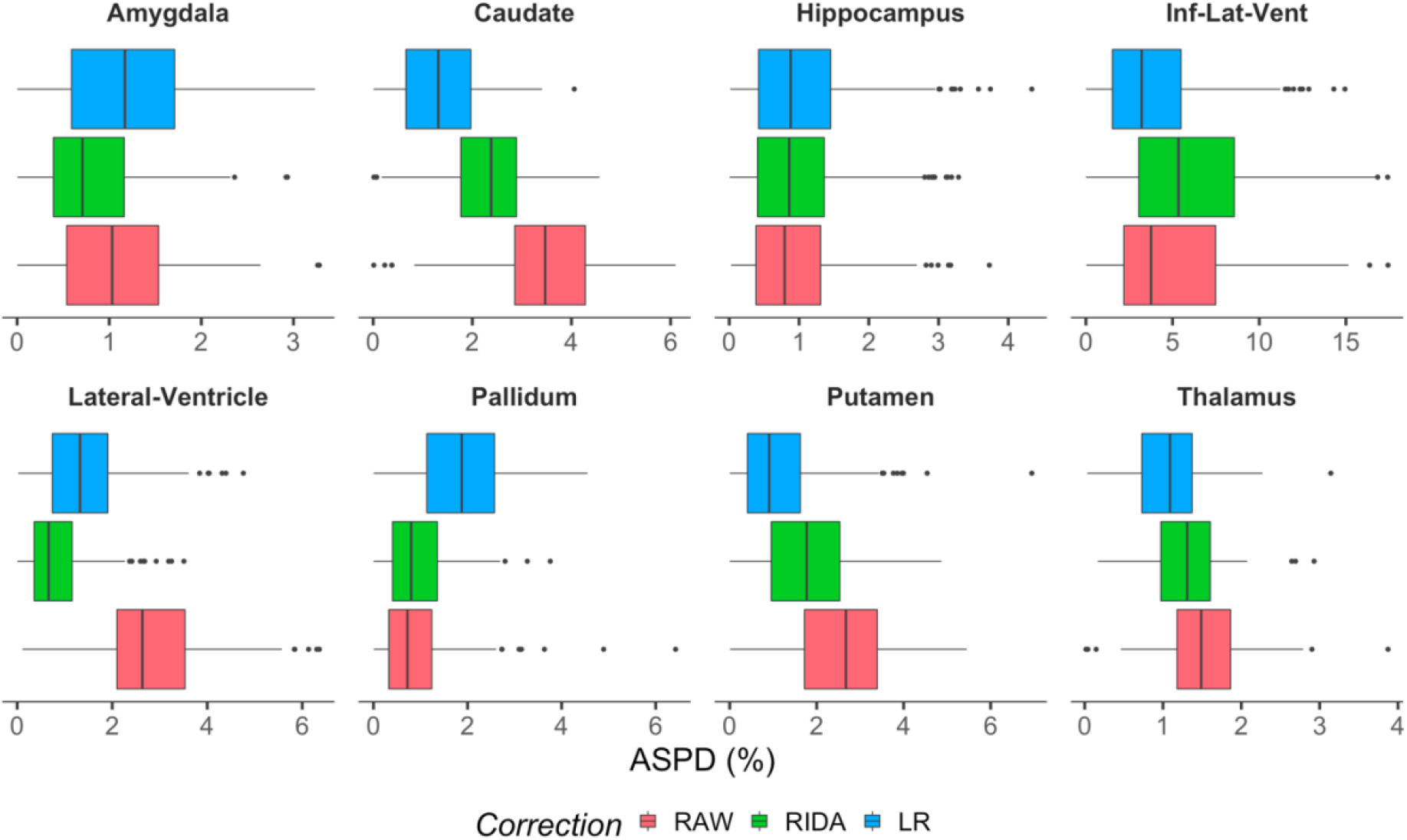
Box plots of within-visit volume absolute symmetrized percentage differences (ASPD) for 243 participants. RAW corresponds to volume differences influenced by the between-scan effects, whereas RIDA (robust intensity distribution alignment) and LR (linear regression) represent volume differences after between-scan effects normalization.

### 3.3. Between-visit longitudinal volume changes

We compared mean ASPCs and standard deviations before and after the between-scan effects correction (Tables 1-3). The results were consistent among age groups – the same brain structures benefited from the same correction methods and thus, did not indicate a bias towards any age group. Volume corrections based on the simple linear regression model yielded more similar longitudinal changes to the reference Avanto-Avanto changes for caudate, inferior lateral ventricle, putamen, and thalamus. The intensity corrections also reduced between-scan effects for the same brain structures but not to the same extent as volume corrections.

**Table 1.** Mean ASPCs and standard deviations (in parenthesis) for the age group of 10 to 20 years. Longitudinal changes: A→A – Avanto-Avanto, A→S – Avanto-Skyra, A→A_pred VC – Avanto-Avanto-predicted after volume corrections, A→A_pred IC – Avanto-Avanto-predicted after intensity corrections. Bolded values indicate volume changes closest to the reference A→A changes. The underlined bolded values indicate significantly different values from A→S changes (p < 0.05).

|  | A→A | A→S | A→ A_pred VC | A→ A_pred IC |
| --- | --- | --- | --- | --- |
| Amygdala | 0.19 (0.35) | 0.38 (0.37) | 0.41 (0.39) | **<u>0.29 (0.36)</u>** |
| Caudate | -0.15 (0.54) | 0.65 (0.49) | **<u>0.16 (0.51)</u>** | 0.38 (0.48) |
| Hippocampus | 0.13 (0.38) | **0.24 (0.40)** | 0.28 (0.41) | 0.25 (0.41) |
| Inf. lateral ventricle | 0.72 (1.93) | 1.55 (2.05) | **<u>0.49 (2.11)</u>** | -0.44 (2.04) |
| Lateral ventricle | 1.12 (1.38) | 1.88 (1.33) | 1.49 (1.35) | **<u>1.26 (1.35)</u>** |
| Pallidum | 0.56 (0.64) | 0.33 (0.62) | 0.87 (0.60) | **<u>0.62 (0.61)</u>** |
| Putamen | -0.25 (0.35) | 0.43 (0.36) | **<u>-0.10 (0.38)</u>** | 0.21 (0.36) |
| Thalamus | 0.01 (0.39) | 0.36 (0.39) | **<u>0.23 (0.39)</u>** | 0.27 (0.39) |

**Table 2.**
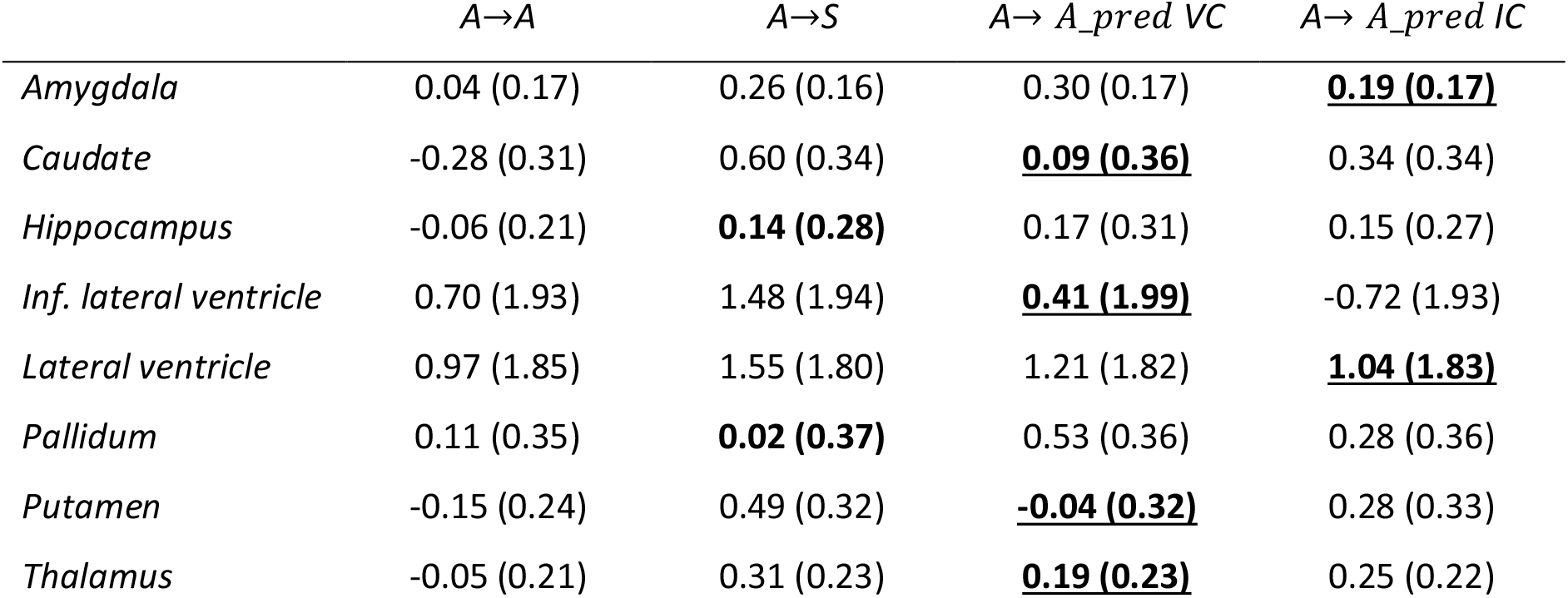
Mean ASPCs and standard deviations for the age group 20 to 40 years. See Table 2 for abbreviations.

|  | A→A | A→S | A→ A_pred VC | A→ A_pred IC |
| --- | --- | --- | --- | --- |
| Amygdala | 0.04 (0.17) | 0.26 (0.16) | 0.30 (0.17) | **<u>0.19 (0.17)</u>** |
| Caudate | -0.28 (0.31) | 0.60 (0.34) | **<u>0.09 (0.36)</u>** | 0.34 (0.34) |
| Hippocampus | -0.06 (0.21) | **0.14 (0.28)** | 0.17 (0.31) | 0.15 (0.27) |
| Inf. lateral ventricle | 0.70 (1.93) | 1.48 (1.94) | **<u>0.41 (1.99)</u>** | -0.72 (1.93) |
| Lateral ventricle | 0.97 (1.85) | 1.55 (1.80) | 1.21 (1.82) | **<u>1.04 (1.83)</u>** |
| Pallidum | 0.11 (0.35) | **0.02 (0.37)** | 0.53 (0.36) | 0.28 (0.36) |
| Putamen | -0.15 (0.24) | 0.49 (0.32) | **<u>-0.04 (0.32)</u>** | 0.28 (0.33) |
| Thalamus | -0.05 (0.21) | 0.31 (0.23) | **<u>0.19 (0.23)</u>** | 0.25 (0.22) |

**Table 3.**
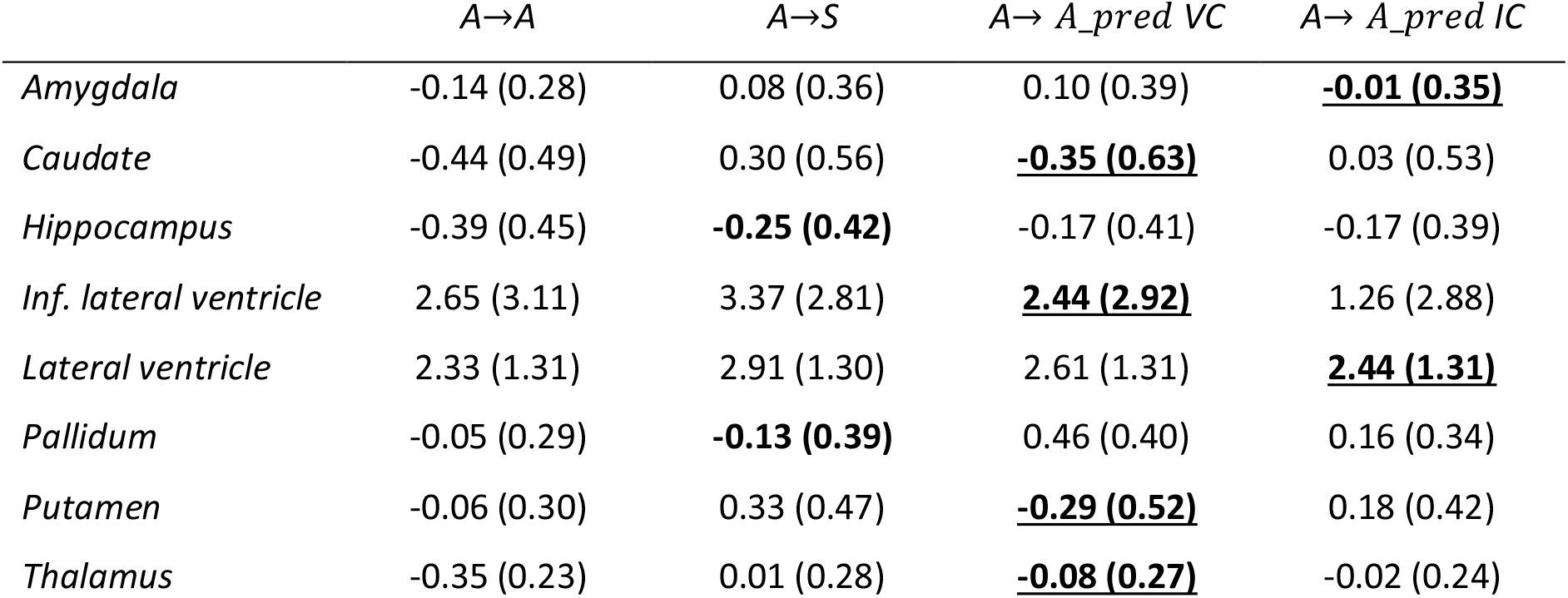
Mean ASPCs and standard deviations for the age group 40 to 81 years. See Table 2 for abbreviations.

|  | A→A | A→S | A→ A_pred VC | A→ A_pred IC |
| --- | --- | --- | --- | --- |
| Amygdala | -0.14 (0.28) | 0.08 (0.36) | 0.10 (0.39) | **<u>-0.01 (0.35)</u>** |
| Caudate | -0.44 (0.49) | 0.30 (0.56) | **<u>-0.35 (0.63)</u>** | 0.03 (0.53) |
| Hippocampus | -0.39 (0.45) | **-0.25 (0.42)** | -0.17 (0.41) | -0.17 (0.39) |
| Inf. lateral ventricle | 2.65 (3.11) | 3.37 (2.81) | **<u>2.44 (2.92)</u>** | 1.26 (2.88) |
| Lateral ventricle | 2.33 (1.31) | 2.91 (1.30) | 2.61 (1.31) | **<u>2.44 (1.31)</u>** |
| Pallidum | -0.05 (0.29) | **-0.13 (0.39)** | 0.46 (0.40) | 0.16 (0.34) |
| Putamen | -0.06 (0.30) | 0.33 (0.47) | **<u>-0.29 (0.52)</u>** | 0.18 (0.42) |
| Thalamus | -0.35 (0.23) | 0.01 (0.28) | **<u>-0.08 (0.27)</u>** | -0.02 (0.24) |

Despite volume corrections indicating more considerable improvements for the caudate, inferior lateral ventricle, putamen, and thalamus, all structures except the hippocampus and inferior lateral ventricles benefited from the intensity correction as well. This was especially evident in the amygdala and lateral ventricles, which showed superior improvements. However, neither of the correction methods yielded better results for the hippocampus and pallidum. These results are in line with the previously communicated within-visit volume changes.

The results of the intensity correction can be related to (Sederevicius et al., 2022), where an exact intensity transformation for each subject was used for the same training dataset. From there, we could have expected reduced between-scan effects for putamen, thalamus, lateral ventricles, and caudate, which was the case in the longitudinal case here as well. Furthermore, the results of another scanner/sequence comparison, for example, Prisma vs. Prisma or Avanto vs. Prisma, indicated that we could expect even more considerable improvements than the Avanto vs. Skyra comparison in this study. Therefore, it can be that this comparison does not yield the best results in all cases for the intensity correction but might be more beneficial in other studies. We also tried a combination of both correction methods: the intensity correction followed by the volume correction. However, we did not find such a setup significantly better than individual approaches.

## 4. Conclusions

The intensity correction (RIDA method) reduced between-scan effects between 1.5T Avanto and 3T Skyra scanners and improved the accuracy of tracking longitudinal volume changes towards the reference comparison (Avanto-Avanto) for all brain structures of interest except the inferior lateral ventricle and hippocampus. This was especially true for the amygdala and lateral ventricles. The statistical volume-based corrections yielded better results for caudate, putamen, and thalamus. However, intensity corrections also improved the accuracy, although not to the same extent as volume corrections. Little variability between subject intensity mappings indicated a reliable CDF-based approach for image harmonization. However, it is hard to say in advance whether the method will benefit a particular study without examining it beforehand. Therefore, a pilot study is needed to investigate the possible advantages of the image harmonization by CDF alignment.

## Acknowledgments

The present research was funded by a grant from Helse-Sør Øst (grant number 2018009), the European Research Council under grant agreements 283634, 725025 (to A.M.F.), 313440 (to K.B.W.), as well as the Norwegian Research Council (to A.M.F., K.B.W.).

## Data and Code Availability

The raw LCBC MRI data supporting the results of the current study may be available upon reasonable request, given appropriate ethical data protection approvals and data sharing agreements. Requests for the raw MRI data can be submitted to the last author Anders M Fjell. Fully-open raw data availability is restricted as participants have not consented to share their data publicly. All data preprocessing and analysis code will be available at https://github.com/LCBC-UiO upon acceptance of the manuscript.

## Appendix A

**Table A. 1.** Numerical results (mean and standard deviation) of within-visit volume absolute symmetrized percentage differences (ASPD) for 243 participants. RAW corresponds to volume differences influenced by the between-scan effects, whereas RIDA (robust intensity distribution alignment) and LR (linear regression) represent volume differences after between-scan effects normalization. Bolded values indicate the lowest mean ASPD for each brain structure.

|  | RAW | RIDA | LR |
| --- | --- | --- | --- |
| Amygdala | 1.05 (0.68) | **0.83 (0.59)** | 1.19 (0.73) |
| Caudate | 3.48 (1.07) | 2.32 (0.89) | **1.37 (0.85)** |
| Hippocampus | **0.92 (0.70)** | 0.99 (0.74) | 1.03 (0.78) |
| Inf. Lateral Ventricle | 4.89 (3.61) | 6.21 (4.08) | **3.91 (3.15)** |
| Lateral Ventricle | 2.83 (1.16) | **0.82 (0.65)** | 1.41 (0.92) |
| Pallidum | **0.93 (0.83)** | 0.93 (0.67) | 1.88 (0.97) |
| Putamen | 2.55 (1.19) | 1.75 (1.03) | **1.18 (1.01)** |
| Thalamus | 1.54 (0.53) | 1.29 (0.43) | **1.07 (0.49)** |

## Literature

Cerri, S., Hoopes, A., Greve, D.N., Mühlau, M., Van Leemput, K., 2020. A Longitudinal Method for Simultaneous Whole-Brain and Lesion Segmentation in Multiple Sclerosis. 2008.05117 [cs, eess] 12449, 119–128. https://doi.org/10.1007/978-3-030-66843-3_12

Chepkoech, J.-L., Walhovd, K.B., Grydeland, H., Fjell, A.M., for the Alzheimer’s Disease Neuroimaging Initiative, 2016. Effects of change in FreeSurfer version on classification accuracy of patients with Alzheimer’s disease and mild cognitive impairment: Effects of Change in FreeSurfer Version. Hum. Brain Mapp. 37, 1831–1841. https://doi.org/10.1002/hbm.23139

Fjell, A.M., Walhovd, K.B., Fennema-Notestine, C., McEvoy, L.K., Hagler, D.J., Holland, D., Brewer, J.B., Dale, A.M., Alzheimer’s Disease Neuroimaging Initiative, 2010. CSF biomarkers in prediction of cerebral and clinical change in mild cognitive impairment and Alzheimer’s disease. J Neurosci 30, 2088–2101. https://doi.org/10.1523/JNEUROSCI.3785-09.2010

Fortin, J.-P., Sweeney, E.M., Muschelli, J., Crainiceanu, C.M., Shinohara, R.T., 2016. Removing inter-subject technical variability in magnetic resonance imaging studies. Neuroimage 132, 198–212. https://doi.org/10.1016/j.neuroimage.2016.02.036

Holland, D., Brewer, J.B., Hagler, D.J., Fennema-Notestine, C., Fenema-Notestine, C., Dale, A.M., Alzheimer’s Disease Neuroimaging Initiative, 2009. Subregional neuroanatomical change as a biomarker for Alzheimer’s disease. Proc Natl Acad Sci U S A 106, 20954–20959. https://doi.org/10.1073/pnas.0906053106

Iglesias, J.E., Van Leemput, K., Augustinack, J., Insausti, R., Fischl, B., Reuter, M., 2016. Bayesian longitudinal segmentation of hippocampal substructures in brain MRI using subject-specific atlases. NeuroImage 141, 542–555. https://doi.org/10.1016/j.neuroimage.2016.07.020

Jack, C.R., Barnes, J., Bernstein, M.A., Borowski, B.J., Brewer, J., Clegg, S., Dale, A.M., Carmichael, O., Ching, C., DeCarli, C., Desikan, R.S., Fennema-Notestine, C., Fjell, A.M., Fletcher, E., Fox, N.C., Gunter, J., Gutman, B.A., Holland, D., Hua, X., Insel, P., Kantarci, K., Killiany, R.J., Krueger, G., Leung, K.K., Mackin, S., Maillard, P., Malone, I.B., Mattsson, N., McEvoy, L., Modat, M., Mueller, S., Nosheny, R., Ourselin, S., Schuff, N., Senjem, M.L., Simonson, A., Thompson, P.M., Rettmann, D., Vemuri, P., Walhovd, K., Zhao, Y., Zuk, S., Weiner, M., 2015. Magnetic resonance imaging in Alzheimer’s Disease Neuroimaging Initiative 2. Alzheimer’s & Dementia 11, 740–756. https://doi.org/10.1016/j.jalz.2015.05.002

Jack, C.R., Knopman, D.S., Jagust, W.J., Petersen, R.C., Weiner, M.W., Aisen, P.S., Shaw, L.M., Vemuri, P., Wiste, H.J., Weigand, S.D., Lesnick, T.G., Pankratz, V.S., Donohue, M.C., Trojanowski, J.Q., 2013. Tracking pathophysiological processes in Alzheimer’s disease: an updated hypothetical model of dynamic biomarkers. Lancet Neurol 12, 207–216. https://doi.org/10.1016/S1474-4422(12)70291-0

Jack, C.R., Knopman, D.S., Jagust, W.J., Shaw, L.M., Aisen, P.S., Weiner, M.W., Petersen, R.C., Trojanowski, J.Q., 2010. Hypothetical model of dynamic biomarkers of the Alzheimer’s pathological cascade. Lancet Neurol 9, 119. https://doi.org/10.1016/S1474-4422(09)70299-6

Jovicich, J., Czanner, S., Greve, D., Haley, E., van der Kouwe, A., Gollub, R., Kennedy, D., Schmitt, F., Brown, G., MacFall, J., Fischl, B., Dale, A., 2006. Reliability in multi-site structural MRI studies: Effects of gradient non-linearity correction on phantom and human data. NeuroImage 30, 436–443. https://doi.org/10.1016/j.neuroimage.2005.09.046

Kassambara, A., 2020. ggpubr: “ggplot2” Based Publication Ready Plots.

Puonti, O., Iglesias, J.E., Van Leemput, K., 2016. Fast and sequence-adaptive whole-brain segmentation using parametric Bayesian modeling. NeuroImage 143, 235–249. https://doi.org/10.1016/j.neuroimage.2016.09.011

R Core Team, 2020. R: A Language and Environment for Statistical Computing. R Foundation for Statistical Computing, Vienna, Austria.

Sederevicius, D., Bjornerud, A., Walhovd, K.B., Leemput, K.V., Fischl, B., Fjell, A.M., 2022. A robust intensity distribution alignment for harmonization of T1w intensity values. bioRxiv 2022.06.15.496227. https://doi.org/10.1101/2022.06.15.496227

Shinohara, R.T., Sweeney, E.M., Goldsmith, J., Shiee, N., Mateen, F.J., Calabresi, P.A., Jarso, S., Pham, D.L., Reich, D.S., Crainiceanu, C.M., 2014. Statistical normalization techniques for magnetic resonance imaging. NeuroImage: Clinical 6, 9–19. https://doi.org/10.1016/j.nicl.2014.08.008

Wickham, H., 2016. ggplot2: Elegant Graphics for Data Analysis. Springer-Verlag New York.

Wickham, H., François, R., Henry, L., Müller, K., 2020. dplyr: A Grammar of Data Manipulation.

Wrobel, J., Martin, M.L., Bakshi, R., Calabresi, P.A., Elliot, M., Roalf, D., Gur, R.C., Gur, R.E., Henry, R.G., Nair, G., Oh, J., Papinutto, N., Pelletier, D., Reich, D.S., Rooney, W.D., Satterthwaite, T.D., Stern, W., Prabhakaran, K., Sicotte, N.L., Shinohara, R.T., Goldsmith, J., 2020. Intensity warping for multisite MRI harmonization. NeuroImage 223, 117242. https://doi.org/10.1016/j.neuroimage.2020.117242

